# Environmental drivers of ranavirus in free-living amphibians in constructed ponds

**DOI:** 10.1101/321299

**Authors:** T.E. Youker-Smith, P.H. Boersch-Supan, C.M. Whipps, S.J. Ryan

## Abstract

Amphibian ranaviruses occur globally, but we are only beginning to understand mechanisms for emergence. Ranaviruses are aquatic pathogens which can cause > 90% mortality in larvae of many aquatic-breeding amphibians, making them important focal host taxa. Host susceptibilities and virulence of ranaviruses have been studied extensively in controlled laboratory settings, but research is needed to identify drivers of infection in natural environments. Constructed ponds, essential components of wetland restoration, have been associated with higher ranavirus prevalence than natural ponds, posing a conundrum for conservation efforts, and emphasizing the need to understand potential drivers. In this study, we analyzed four years of *Frog virus 3* prevalence and associated environmental parameters in populations of wood frogs (*Lithobates sylvaticus*) and green frogs (*Lithobates clamitans*) in a constructed pond system. High prevalence was best predicted by low temperature, high host density, low zooplankton concentrations, and Gosner stages approaching metamorphosis. This study identified important variables to measure in assessments of ranaviral infection risk in newly constructed ponds, including effects of zooplankton, which have not been previously quantified in natural settings. Examining factors mediating diseases in natural environments, particularly in managed conservation settings, is important to both validate laboratory findings *in situ*, and to inform future conservation planning, particularly in the context of adaptive management.

## Introduction

Ranaviruses are primarily infectious pathogens of aquatic ectothermic vertebrates, and have been implicated in mass die-offs of amphibians worldwide (Duffus et al. 2015). Many anuran species, including the wood frog (*Lithobates sylvaticus*), experience nearly 100% mortality after exposure as larvae (Hoverman et al. 2011). Little is currently known about reasons for ranavirus emergence, although anthropogenic disturbance is suspected as a leading factor (Jancovich et al. 2005; Forson and Storfer 2006; Storfer et al. 2007; Miller et al. 2009). With increasing awareness of potential human influence, research efforts are aimed at identifying risk factors for infection with the goal of reducing both spread and persistence of ranaviruses. In this study we identified factors influencing prevalence of *Frog virus 3* (FV3), a widespread ranavirus, in populations of *L. sylvaticus* and green frogs (*Lithobates clamitans*) in a constructed vernal pool array in New York State, United States, by developing statistical models of prevalence in response to environmental and organism-focused variables.

Individual factors have been evaluated in controlled settings to determine virulence of ranaviruses both in the environment and within hosts. Ranaviral replication rates *in vitro* generally increase as temperature increases (Ariel et al. 2009); however, ranaviral infectivity declines at a faster rate at higher temperatures (Nazir et al. 2012; Munro et al. 2016). Ranaviruses may persist in the environment for several weeks to months in dry conditions (Nazir et al. 2012; Munro et al. 2016). This raises concern when examining recurring outbreaks because many wetland types that support populations of aquatic-breeding anurans in the northeastern United States may partially or completely dry up during late summer or over winter. Furthermore, the interplay of pond-drying and other abiotic factors on the prevalence and infection dynamics of aquatic diseases remains poorly understood (Paull and Johnson 2018). Vernal pools also support highly diverse microbial and micro-invertebrate communities, and although less rigorously studied, these communities could be highly influential in understanding outbreak etiology. For example, FV3 becomes less virulent in the presence of zooplankton (Johnson and Brunner 2014), and survives longer in filtered and sterilized water (Nazir et al. 2012; Johnson and Brunner 2014; Munro et al. 2016).

In addition to environmental conditions, we examined several variables shown to affect susceptibility of amphibians to FV3 including developmental stage, and density. Although both *L. sylvaticus* and *L. clamitans* have relatively high probabilities of infection and mortality when exposed to FV3, we expect highest prevalence rates overall in *L. sylvaticus* (Hoverman et al. 2011). Water temperature produces different results with respect to infectivity and mortality, depending on both *Ranavirus* strain and host species (Rojas et al. 2005; Bayley et al. 2013; Echaubard et al. 2014; Brand et al. 2016). In regards to specifically FV3 and anuran ranid species (which includes *L. sylvaticus* and *L. clamitans*), research has produced conflicting results. Many controlled studies supported a positive correlation, with higher mortality rates at warmer temperatures (Bayley et al. 2013; Brand et al. 2016). In contrast, Echaubard et al. (2014) and Gray et al. (2007) found that probability of both infection and mortality was *lower* at warmer temperatures. In a natural setting, seasonal increases in temperature generally correspond with progression towards metamorphosis in aquatic anuran larvae, measured by increases in Gosner developmental stage (Gosner 1960). When examining Gosner stage alone, different species exhibit differing trends in susceptibility, but in ranid species infection and mortality generally increase as larvae approach metamorphosis (Haislip et al. 2011; Warne et al. 2011). It is unclear what role host density may play in FV3 outbreaks, as response to density is non-linear; other factors such as behavior, metamorphic rates, and baseline host fitness differ in low versus high density conditions and blur the effects of ranaviruses (Greer et al. 2008; Echaubard et al. 2010; Reeve et al. 2013).

Over a four-year study period, we recorded estimated FV3 prevalence and developed explanatory models of prevalence in response to temperature, larval density, Gosner stage, spatial clustering of pools, and zooplankton communities. The objectives of this study were to better understand the influence of environmental and host conditions on FV3 outbreaks in natural settings, furthermore we specifically wanted to quantify the effect of zooplankton on FV3 prevalence within natural systems. The use of newly constructed ponds in the study site presented a unique opportunity to assess FV3 risk in ponds that have a known history and were monitored since their creation.

## Methods

### Study site

Svend O. Heiberg Memorial Forest (42° 46’ N, 76° 5’ W), is a 1,600 ha property owned and maintained by the State University of New York College of Environmental Science and Forestry (SUNY ESF). An array of 71 vernal pool basins (Figure 1A) was constructed in 2010 by SUNY ESF and the Upper Susquehanna Coalition, to recreate *L. sylvaticus* and spotted salamander (*Ambystoma maculatum*) breeding habitat previously destroyed by land use change associated with the sequence of forest clearance, intensive agriculture, and subsequent agricultural abandonment and forest regrowth over the last two centuries. Pools varied from 3m-10m diameter, with most circular or ovular in shape. Pools were designed to be hydrologically isolated and were arranged in clusters of 1, 3, or 9 pools within 164m-diameter landscape hexagons (Figure 1C). A separate cluster of 32 pools, the “microarray” (Figure 1B), was constructed in a grid pattern spanning forested, field, and edge habitats. Several naturally occurring vernal pools were also present within hexagon clusters.

**Figure 1:**
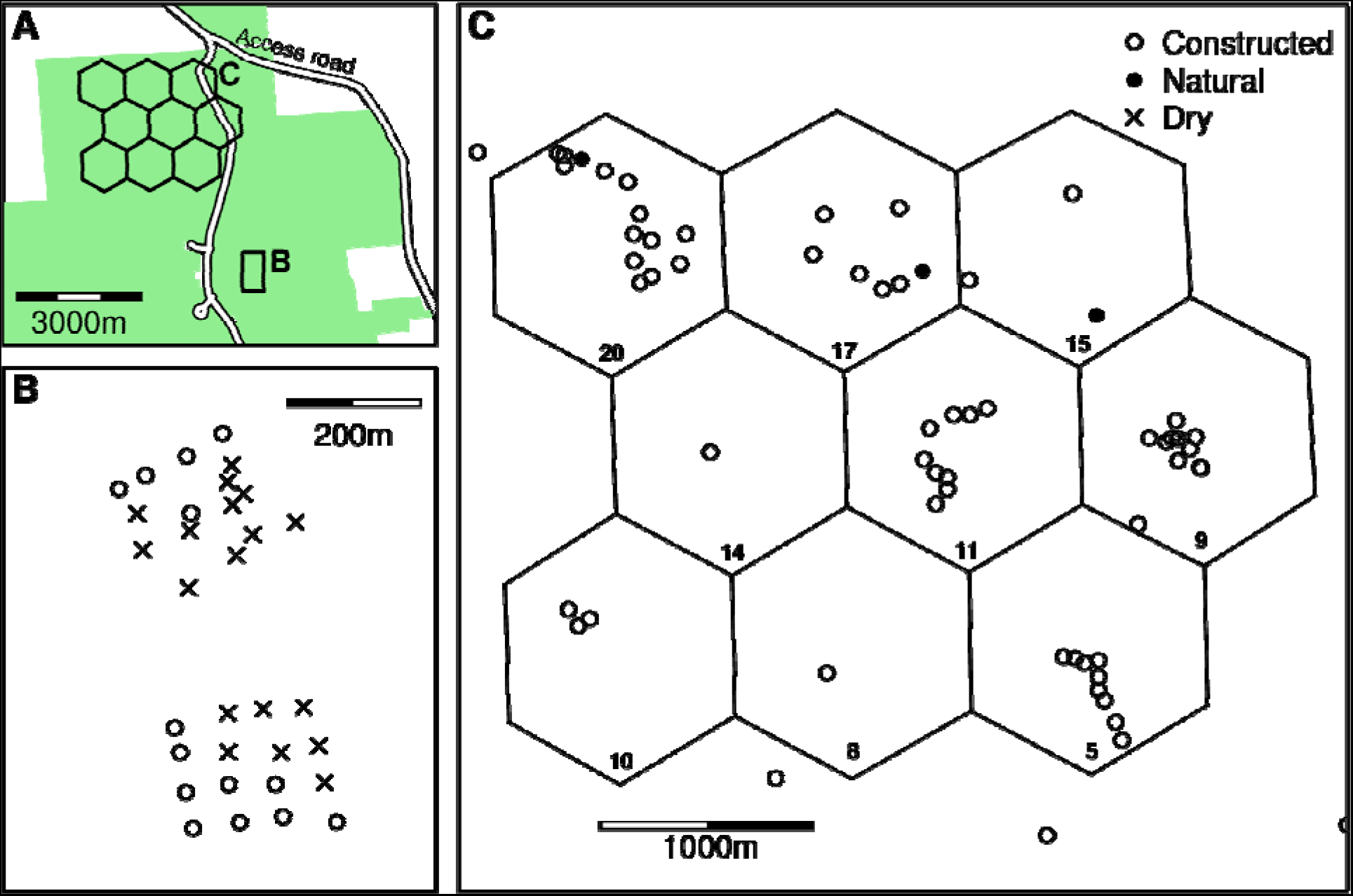
(A) Vernal pool array within Heiberg Memorial Forest (shaded area). Pools were constructed (B) in a separate grid- patterned microarray, as well as (C) in clusters of 1, 3, or 9 pools within uniform landscape hexagonal areas. Study pools included constructed pools (open circles) with three pre-existing pools (filled circles). Sixteen pools failed to hold water at any point in the study, here designated as “dry” (crosses).

### Sampling

All constructed ponds containing water and four natural ponds were sampled at three separate intervals during *L. sylvaticus* larval development from 2011–2014. Sampling events were spaced three to four weeks apart and began approximately six to eight weeks after *L. sylvaticus* egg masses were observed, allowing tadpoles to develop to at least Gosner stage 25 (Gosner 1960). First sampling intervals occurred from mid-May to early June, depending on timing of spring thaw and *L. sylvaticus* breeding events. Sampling in 2013 was restricted to one interval in June-July.

Larval sampling at each interval was performed by modified pipe sampling methods as described in Werner et al. (2007). A 33 cm-diameter section of spiral duct pipe was plunged through the water column into the sediment, and tadpoles trapped within the pipe were collected by net sweeps and stored in buckets with water from the same pool. A sample was considered empty once zero individuals were captured for ten consecutive net sweeps. Samples were spaced at least 2 m apart with the exception of pools less than 5 m, from which approximately one sample per 2 m^2^ of surface area was taken. Equipment was immersed in 10% bleach solution for at least 60 seconds and allowed to air dry between pools. Thirty tadpoles were randomly selected for processing in pools where at least 30 were captured, and all tadpoles were used in pools where less than 30 were captured. All other individuals were immediately returned to their pool of origin. Selected individuals were humanely euthanized by immersion in 70% ethanol, and stored in 95% ethanol at 4°C until further processing. Sampling was performed according to State University of New York College of Environmental Science and Forestry IACUC protocol #140201.

### Environmental and organism-focused parameters

Marked wooden stakes were driven into the sediment in the estimated deepest area of each pool, and visited weekly to record water depth from the first spring thaw until November. Two temperature loggers (Thermochron^®^ iButtons^®^, Embedded Data Systems, Lawrenceburg, KY) per pool were attached to 15 cm lengths of copper wire and coated with Performix^®^ Plasti Dip^®^ (Plasti Dip International, Blaine, MN). A length of epoxy coated rebar greater than the maximum depth for each pool was driven into the sediment near depth stakes, and one thermal logger was attached at the bottom of each pool. One thermal logger was affixed to the bottom of a foam float attached loosely to freely move up and down posts, to measure surface temperature. Thermal loggers were programed to record readings every three hours, and were retrieved and redeployed every six months during the study period. Zooplankton concentrations were taken from Holmes et al. (2016). Briefly, zooplankton were sampled by passing 3 L of water, taken from the center of each pond, through an 80-µm sieve. Animals were preserved in 95% ethanol and manually counted and identified to the species level, or lowest taxonomic group possible when species could not be identified (Holmes et al. 2016).

### Ranaviral DNA screening methods

#### PCR assays

Screening for the presence of ranaviral DNA followed the methods outlined in Youker-Smith et al. (2016), and a full description of the employed protocols is given in the supplementary materials. Briefly, DNA was isolated and purified from up to 25 mg tadpole liver tissue using a modified salt extraction method (Sambrook and Russell 2001). Template DNA (5 µL) was then amplified with conventional PCR using primers for *Frog virus 3* major capsid protein (MCP) 4 and 5 as described in Mao et al. (1997; MCP 4: 5′-GAC TTG GCC ACT TAT GAC-3′; MCP 5: 5′-GTC TCT GGA GAA GAA GAA-3′). Amplified base pair segments were separated by 1% agarose gel electrophoresis and stained with ethidium bromide for visualization. Sequenced DNA from a dead *L. sylvaticus* tadpole sampled in June 2011 from Hexagon 11 was used as positive control. Negative and ambiguous results were re-amplified using the above methods to increase screening sensitivity. For the 2014 samples negative or ambiguous results were re-analyzed via quantitative PCR using a protocol modified from Pallister et al. (2007).

### Sequencing

Amplified PCR products from four dead or moribund *L. sylvaticus* collected from die-offs in Hexagons 11 and 5 (Figure 1C) in 2011 were purified using Omega E.Z.N.A.^®^ Cycle Pure Kit (Omega Bio-tek Inc., Norcross, GA). Purified products were sequenced at the Yale University DNA Analysis Facility. Sequences were aligned with BioEdit v 7.2.5 and a GenBank (Clark et al. 2016) sequence search performed using nucleotide BLAST^®^, targeting nucleotide collection entries optimized for highly similar sequences (Johnson et al. 2008).

### Data Analysis

*Frog virus 3* (FV3) prevalence was modeled using hierarchical generalized linear regression models (GLMs) with binomial error distribution and logit link function in response to the following variables: temperature, water depth, Gosner developmental stage, tadpole host density, water depth, the average distance to the three nearest neighboring pools (as a measure of spatial clustering of pools), and total pelagic zooplankton concentration (Table 1). No temperature or zooplankton data were available for the single sampling interval in 2013, and the corresponding prevalence data were therefore excluded from the statistical analysis.

**Table 1.**
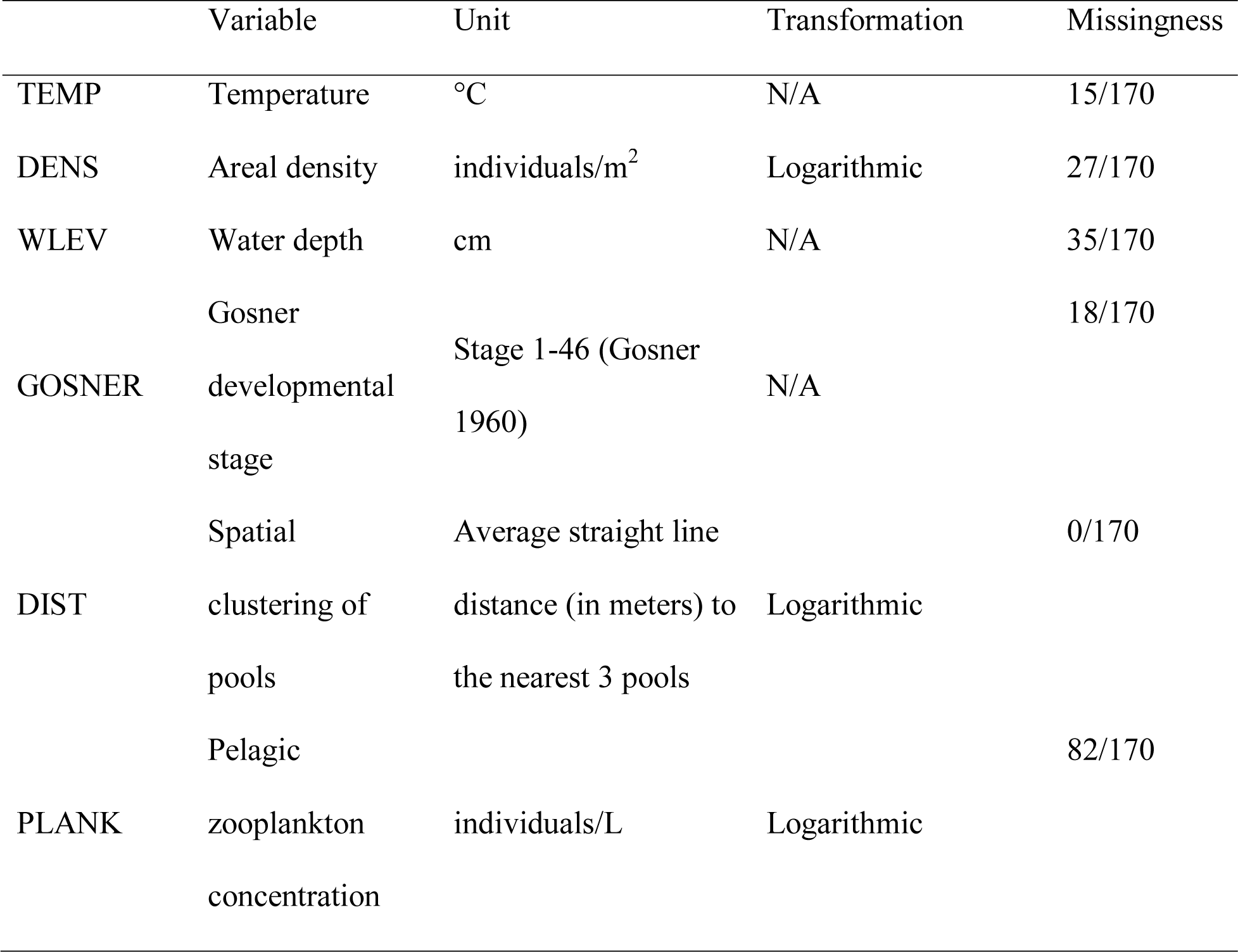
List of explanatory variables for generalized linear models of Frog virus 3 (FV3) prevalence in vernal ponds. Missingness gives the proportion of FV3 prevalence observations for which no corresponding observation of a particular environmental covariate was available.

**Table 2.** Posterior estimates of regression coefficients for the best predictive model

|  | Mean | StdDev | 95% credible interval |  |
| --- | --- | --- | --- | --- |
| Intercept | 3.22 | 2.37 | -1.30 | 7.60 |
| GOSNER | 0.08 | 0.05 | -0.01 | 0.17 |
| TEMP | -0.25 | 0.07 | -0.40 | -0.11 |
| log(DENS) | 0.32 | 0.13 | 0.08 | 0.58 |
| WLEV | 0.05 | 0.01 | 0.03 | 0.07 |
| log(PLANK) | -0.63 | 0.06 | -0.75 | -0.51 |
| log(DIST) | -0.70 | 0.17 | -1.04 | -0.37 |
| sigma_W | 1.54 | 0.49 | 0.84 | 2.63 |

GLM parameters were estimated in a hierarchical Bayesian framework using the rstan and rethinking packages in R. This inference framework provided a coherent approach to modelling missing predictor values, which was essential to maintain a dataset representative of the sampling design, given that zooplankton data were only available for approximately 50% of samples (Table 1). Apart from the interpond distances, all other predictor variables exhibited a lesser degree of missingness (Temperature 9%, Host density 16%, Water level 20%, Gosner stage 10%) as a result of logistical constraints on sampling and/or equipment failures.

A hierarchical model structure was chosen to accommodate so-called random intercepts for each of the nine sampling occasions. Model structure was as follows:

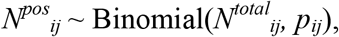

where *N^pos^_ij_* are the number of FV3 positive tadpoles out of a sample of *N^total^_ij_* in pool *i* at sampling occasion *j*, and *p_ij_* is the expected FV3 prevalence, modeled itself as 
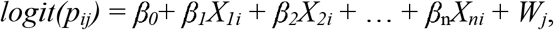
where β_*1…n*_ are the regression coefficients for predictors *X_1…n_* for pool *i*. *W_j_* is the random intercept for sampling interval *j*, and was modelled as *W_j_* ∼ Normal(0, *σ_W_*). Further, missing predictor values *X_ni_* were estimated as coming from a normal distribution *X_ni_* ∼ Normal*(μ_Xn_*, *σ_Xn_*), the mean and variance of which were estimated from the observed values of each predictor jointly with all other model parameters.

Following recommendations in Gelman et al. (2008) and McElreath (2016) we employed weakly informative priors to regularize extreme inferences that can be obtained using maximum likelihood or completely non-informative priors. Normal(0,10) priors were chosen for all regression coefficients, Half-Cauchy(0, 2) priors for all variance parameters. The priors for the mean *μ_Xn_* of a missing predictor values followed a normal distribution centered on the mean of the observed predictor values with a standard deviation of 10.

Candidate models with different predictor combinations were evaluated using leave-one-out cross-validation (LOO) as implemented in the loo package in R (Vehtari et al. 2017), and ranked using the LOO information criterion IC_LOO_. IC_LOO_ is asymptotically equal to the Watanabe-Akaike information criterion (WAIC; Watanabe 2010) as a means for estimating pointwise out-of-sample prediction accuracy, but is more robust for finite sample sizes (Vehtari et al. 2017).

## Results

DNA sequences obtained from four *L. sylvaticus* individuals exhibiting ranavirus pathologies in 2011 shared 100% identity with *Frog virus 3* isolate D1 major capsid protein gene (GenBank accession JQ771299). *Frog virus 3* site-wide prevalence ranged from 0.03 to 0.57.

The model with the highest predictive accuracy included temperature, Gosner stage, water level, host density, zooplankton density and a measure of spatial clustering as predictors. Within this model, FV3 prevalence decreased with water temperature (β = −0.25, 95% CI (−0.40,−0.11); Fig. 2A). and increased with increasing water level (β = 0.05, 95% CI(0.03,0.17); Fig. 2C). Further, prevalence increased with an increase in host density (β = 0.32, 95%CI (0.08, 0.58); Fig. 2D), but decreased markedly with increasing zooplankton densities (β =−0.63, 95%CI (−0.75, –0.51); Fig. 2E). Prevalence decreased slightly with increasing distance to neighboring pools (β = –0.70, 95% CI (−1.04, –0.37); Fig. 2F), although a model without this predictor had a similar predictive accuracy (ΔIC = 8, SE 18; Table S1). There was also some evidence that prevalence increased slightly as frogs approached metamorphosis (i.e. Gosner stage 42; (β = 0.08, 95% CI (−0.01,0.17); Fig. 2B). Models incorporating fewer covariates exhibited a substantially lower predictive accuracy (Table S1).

**Figure 2:**
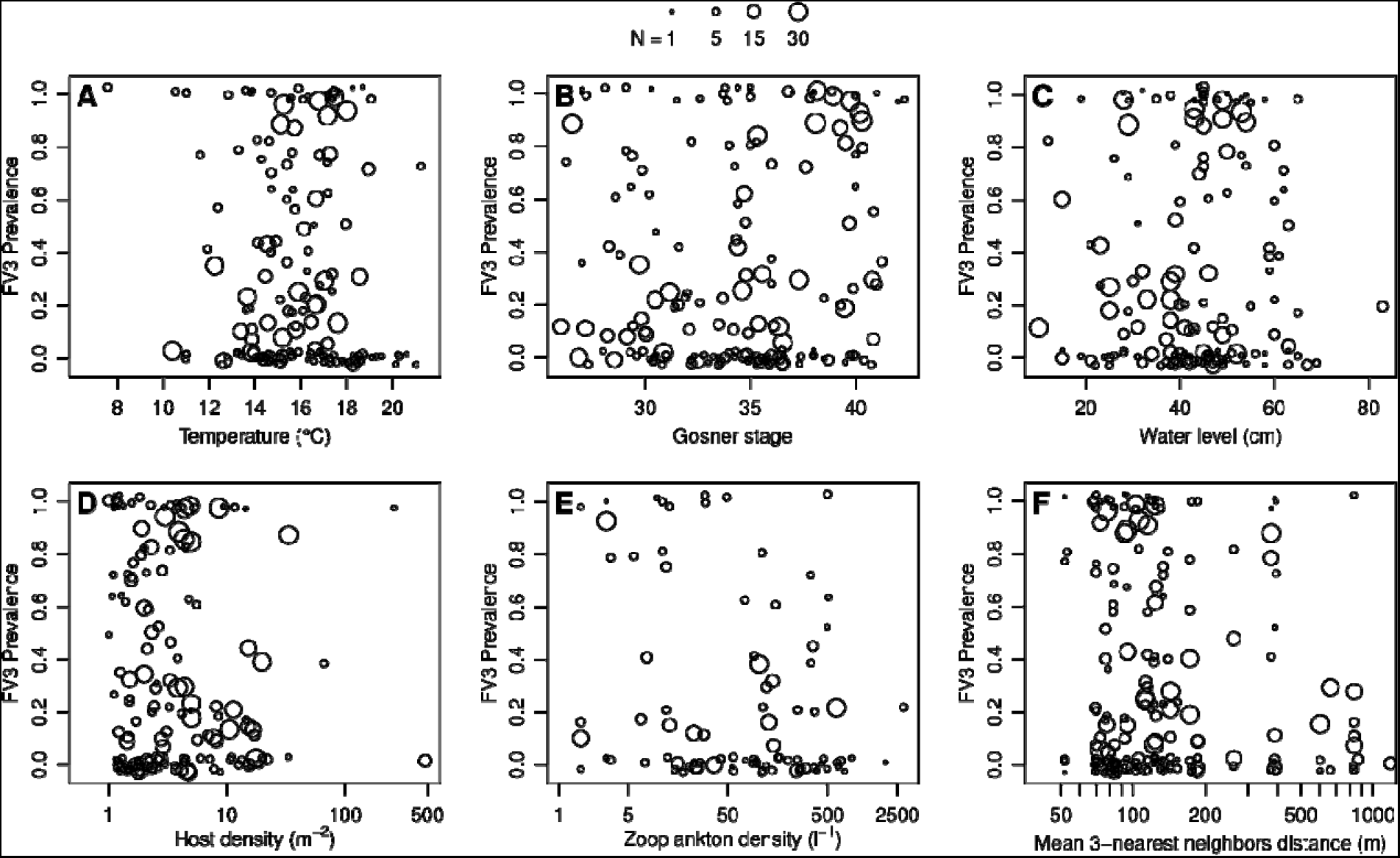
Observed Frog virus 3 prevalence in relation to environmental covariates water temperature (A), developmental stage (B), pond water level (C), host density (D), zooplankton density (E), and average distance to the three neirest neighboring ponds (F) on Frog virus 3 prevalence. Prevalence values are jittered along the y-axis by up to 0.03 units to alleviate overplotting. Symbol size reflects the number of successfully assayed tadpoles in a given sample.

**Figure 3:**
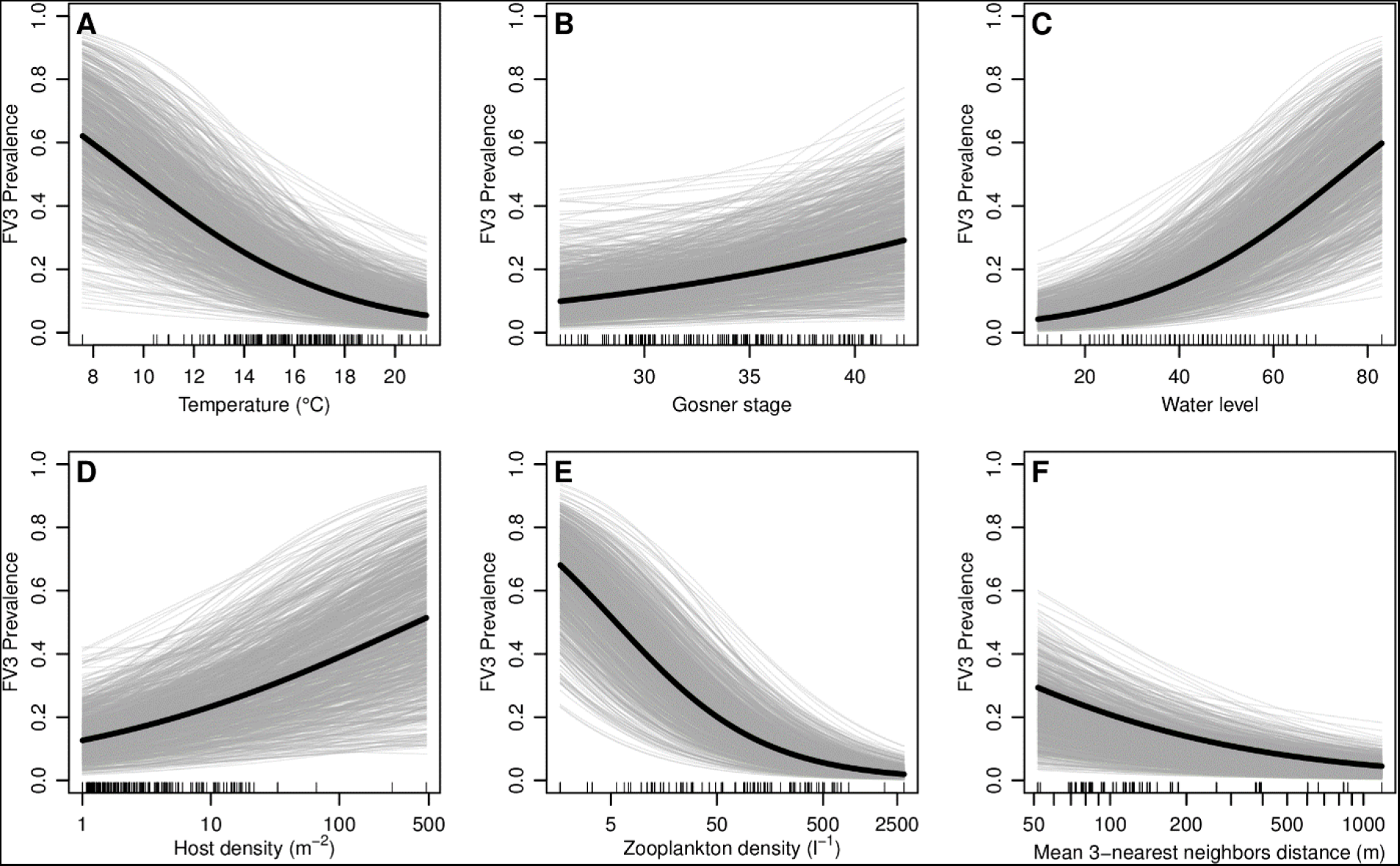
Partial effects of environmental covariates water temperature (A), developmental stage (B), pond water level (C), host density (D), zooplankton density (E), and average distance to the three neirest neighboring ponds (F) on Frog virus 3 prevalence, as estimated in the best predictive model. Black lines represent posterior means of regression coefficients, grey lines are 1000 draws from the posterior of each regression coefficient. Predictor observations are indicated by the black tickmarks along the x-axes.

## Discussion

The results of this study showed that low temperature, high host density, low zooplankton concentrations, deep water, the close vicinity of other pools, and host Gosner stages approaching metamorphosis were predictors of high FV3 prevalence. These results showed how responses of ranaviruses and hosts to environmental conditions tested in controlled laboratory, or even mesocosm, experiments, may not be representative of what can be expected in a natural setting. These findings also provided novel evidence that zooplankton may play a significant role in reducing prevalence of ranaviruses in the natural environment – a phenomenon previously only studied in laboratory settings (Johnson and Brunner 2014).

Individual parameters in this study did not conform to previously reported results for several possible reasons. Water temperature was included in all best fit candidates for both models, and produced negative trends with respect to FV3 prevalence. This was in contrast to controlled studies supporting positive trends (Bayley et al. 2013; Brand et al. 2016). However, unlike in laboratory settings, temperature levels fluctuated with daily and seasonal cycles and were not controlled and/or stable in these wild populations. Temperature, as measured in this study, therefore likely also captures other aspects of the forest environment, and interacts with other variables in the natural environment. Temperature effects are therefore not straightforward to compare between laboratory studies and observations of free-living populations. As previously mentioned, and as demonstrated e.g. in the frog–chytrid fungus system (Raffel et al. 2013), effects of temperature on ranaviruses depend on both host susceptibility/immunity and pathogen replication/virulence. In this study, the detrimental effects of cold temperatures on host immunity may have overshadowed the effects of cold temperatures on FV3 virulence. Future studies at Heiberg should include surveillance of *L. clamitans* tadpoles in the fall, after *L. sylvaticus* have metamorphosed and temperatures decrease.

Although the credible interval for the effect of developmental stage on virus prevalence did contain zero, over 90% of the posterior mass was positive, indicating increased prevalence with Gosner stage, which is generally consistent with other literature regarding certain ranids (Haislip et al. 2011; Warne et al. 2011). However, Gray et al. (2007), in a study of FV3 prevalence in Tennessee wetlands, found no significant trends between prevalence and Gosner stage for *L. clamitans*. Similarly, Haislip et al. (2011) found no difference in infection or mortality rates between larval and metamorph stages in *L. sylvaticus* or *L. clamitans*. Overwintering *L. clamitans* should also be compared to *L. clamitans* who hatch and complete metamorphosis in the same year. Metamorphosis may be quantified by both growth rates (i.e. length and mass) and differentiation rates (i.e., the rate at which larvae progress through each Gosner stage). *Lithobates clamitans* who overwinter continue slow growth but cease differentiation during the coldest winter months (Smith-Gill and Berven 1979), and we do not yet know what effects this may have on probability of FV3 infection or mortality.

Density was included as a predictor in best fit models. Prevalence increased with increasing density, as may be expected from increased contact rates. Density-dependent infection with ranaviruses have been suggested based on some field studies (Green et al. 2002, Brunner et al. 2004, but in other studies was either not a significant factor (Harp and Petranka 2006) or deemed “not a factor” (Gray et al. 2007) due to field observations and knowledge of study species, which we include in our subsequent discussion. Generally, “host density” from a disease transmission perspective is incredibly difficult to assess in a natural setting (and specifically this study) for several reasons. Larval amphibians other than the target species were often present in pools and their density was not quantified. These species often occupied the same feeding niches and aquatic zones (e.g. *A. maculatum* mostly remained in warm littoral zones or under leaf litter – the same areas in which *L. clamitans* most often occurred; *personal observation*), thus potentially contributing to stress and greater transmission rates from increased contact. When overall pool density was low, tadpoles would often aggregate, thus increasing rates of contact. This same phenomenon was observed by Greer et al. (2008), in a study of density and ATV transmission among tiger salamanders (*Ambystoma tigrinum*). It is also virtually impossible in a natural setting to differentiate between density of infected individuals and density of susceptible hosts. Infection and mortality rates largely depend on the viral dose at which susceptible hosts are exposed (Brunner et al. 2005; Echaubard et al. 2010), and susceptible host density alone does not have significant effects on either infection or mortality (Greer et al. 2008; Echaubard et al. 2010; Reeve et al. 2013).

The best fit model also predicted an increase in prevalence with increasing water depth. This may in part be a reflection of the negative temperature-prevalence relationship we found, as deeper pools tend to have lower water temperatures. Furthermore, deeper pools would tend to not completely freeze down to the sediment during the winter, which may support overwintering of infected green frogs.

Non-amphibian community assemblages are often overlooked in studies of ranaviruses, and microbial and microinvertebrate communities may have substantial effects on pathogen virulence. Ranaviruses survive longer in sterilized environments (Nazir et al. 2012; Johnson and Brunner 2014; Munro et al. 2016), suggesting microbial competition may be a factor in reducing replication rates and infectivity. Zooplankton, specifically *Daphnia* spp., have been studied as potential biological control agents for another deadly amphibian disease, chytrid, caused by the fungus *Batrachochytrium dendrobatidis*. *Daphnia* spp. ingest zoospores and significantly decrease concentrations of *B. dendrobatidis* in the environment (Buck et al. 2011). Johnson and Brunner (2014) observed a similar phenomenon with *Daphnia* and FV3; although *Daphnia* did not decrease the *abundance* of FV3, *infectivity* was reduced. The authors speculated virus particles were somehow mechanically inactivated by the digestive processes of *Daphnia*. In this study, *Daphnia* observations were too sparse to use as a predictor, but total zooplankton (which included *Daphnia* spp.) was a predictor that substantially improved predictive accuracy of models (Table S1). Pools with high zooplankton concentrations had substantially lower FV3 prevalence than pools with less than c. 50 individuals per liter (Fig. 2E). This finding suggests microinvertebrate communities may have been overlooked thus far in the field of amphibian ranavirus research. Although *Daphnia* have been previously studied in controlled laboratory experiments, other zooplankton should be included in future research; in this study, “total zooplankton” also included copepods, ostracods, and non-*Daphnia* cladoceran species (Holmes et al. 2016).

Clustering of pools had a small effect on FV3 prevalence, although including this parameter only provided a marginal improvement of predictive accuracy, when all other predictors were also considered. Spatial characteristics should be important drivers of transmission as sub-lethally infected adults travelling between sites could be sources of infection; however, we did not find this to be a strong predictor of FV3 prevalence in this system (also see Gahl and Calhoun 2008; Greer et al. 2009). Other potentially predictive parameters in future studies may be pool geographic locations, as pools at lower elevations and therefore lower catchment areas could receive more inputs from runoff (Gahl and Calhoun 2008).

Surveillance methods were not adequate to make inferences about FV3 transmission dynamics in this system, given the relatively sparse sampling in time, and because logistical constraints prohibited us from sampling all potential sites harbouring outbreaks. Surveillance for FV3 detection in this study included the two most commonly observed anuran species, but in future studies involving transmission, other amphibian taxa must be considered. Several other larvae of aquatic-breeding amphibians were observed in the study pools, including (in order of decreasing abundance) spring peepers (*Pseudacris crucifer*), spotted salamanders (*Ambystoma maculatum*), American toads (*Anaxyrus americanus*), Eastern red-spotted newts (*Notophthalmus viridescens*), and American bullfrogs (*Lithobates catesbeianus*). Each of these species is susceptible, to some degree, to ranaviruses (Green et al. 2002; Gahl and Calhoun 2010; Hoverman et al. 2011; Hoverman et al. 2012; Richter et al. 2013) and could be additional sources of infection for *L. sylvaticus* and *L. clamitans*. Sub-clinically infected adults of these species also serve as potential reservoirs and may introduce ranaviruses to other populations, or re-introduce the virus in subsequent years (Brunner et al. 2004; North et al. 2015).

Pool geomorphology is also an important consideration especially with constructed ponds. Higher prevalence of ranaviruses has been associated with constructed vs natural wetlands, which has been attributed to deeper basin shapes with little to no littoral zones, longer hydroperiods and less emergent vegetation (Petranka et al. 2003; Greer and Collins 2008; Richter et al. 2013), and utilization of ponds for cattle (Gray et al. 2007). Although study ponds at Heiberg were constructed, they were not representative of the “constructed ponds” referenced in the literature as having higher prevalence for several reasons. The Heiberg pools were located within a mainly densely forested landscape with no agricultural use or livestock access. Most pool basins were gradually sloping, creating the broader littoral zones characteristic of natural pools. Although we did not quantify aquatic vegetation, we observed abundant vegetation (submergent, emergent, and free-floating) in many constructed ponds during sampling. Vegetation is a recommended parameter to include in future studies, as tadpoles may be more spatially distributed in ponds with greater vegetation thus decreasing rates of contact (Greer and Collins 2008). In these regards, constructed ponds at Heiberg appeared to mimic natural systems, with the exception of hydroperiod. Most ponds remained permanently filled, and the few that did not either contained no amphibian larvae or dried before larvae could reach metamorphosis. Ranaviruses cause mortality and may lead to reduced fitness, but aquatic breeding amphibians in particular are already subject to an onslaught of challenges prior to metamorphosis, with field mortality rates for larval anurans exceeding an average of 90% (Melvin and Houlahan 2012). This makes it difficult to determine the degree to which ranavirus-caused mortality exceeds the background rate. Continued disease surveillance therefore needs to be coupled with longitudinal population monitoring to detect long-term population effects of ranavirus prevalence, however our study has provided additional insights into ways of immediately reducing ranavirus infection and mortality in newly constructed ponds closely mimicking natural systems. Many factors must be taken into consideration when designing constructed wetlands such as – to name just a few – proximity to anthropogenic influence, hydrological catchment, availability of amphibian source populations, predation risk e.g. accessibility of the wetland to fish. In addition, by designing ponds with locations and basin geomorphologies favoring warmer temperatures, and stocking to establish a plankton community, we may further reduce disease risk and promote thriving populations in artificial wetlands.

## Acknowledgments

We thank the Maurice Alexander memorial research fund, Western New York Herpetological Society, and SUNY ESF Graduate Student Association for funding. We also thank the Upper Susquehanna Coalition and SUNY ESF for access to forest properties, JB for molecular protocols, RO at Cornell University Animal Health Diagnostic Center, and JA, LO, LL, CR, GG and RS for field and laboratory assistance. Collection methods for live amphibians followed approved SUNY ESF IACUC protocol #140201.

## Supplementary Materials for Environmental drivers of ranavirus in free-living amphibians in constructed ponds

### Supplementary Methods - Quantitative PCR protocol

Negative or ambiguous results were analyzed via quantitative PCR at Cornell University Animal Health Diagnostic Center using the following protocol modified from Pallister et al. (2007): Five μL template DNA was added to 5 μL Invitrogen TaqMan^®^ Fast Virus 1-Step Master Mix, 0.05 μL fluorescent probe (100 μM; 5’-CAC AAC ATT ATC CGC ATC-3’), and 0.18 μL primers (100 μM; rtMCP-F: 5’-CTC ATC GTT CTG GCC ATC AA-3’; rtMCP-R: 5’-TCC CAT CGA GCC GTT CA-3’) to a total volume of 20 μL. Samples were run alongside negative and positive controls in 48-well plates using Applied Biosystems StepOne™ real-time PCR system and analyzed with StepOne software v2.3. A synthetic Ultramer^®^ oligomer containing binding sites from primers and probe described above was used as positive control (R. Ossiboff, Cornell University Animal Health Diagnostic Center; 5’- AAG ACT TGG CCA CTT ATG ACT TGC ATC GGC AGC AAA TCT CAT CGT TCT GGC CAT CAA CCA CAA CAT TAT CCG CAT CAT CAA CGG CTC GAT GGG ATG CCA TAT TTT AAG AGA ATT ATC GAG GTC TCT GGA GAA CAA GAA CG – 3’). Five serial dilutions of 1:10 were run in duplicate and used to calibrate a set of standards, and a cycle threshold (C_T_) was set at the logarithmic center of standard linear growth curves for each run. Samples with C_T_ < 36 were declared positive; this threshold was based on mean C_T_ values of the lowest concentration of positive control (1 × 10^3^ nM).

### Supplementary Table – Model selection

**Table S1:** Model selection results for generalized linear models of Frog virus 3 prevalence. IC_LOO_: Leave-one-out cross validation Information Criterion; p_LOO_: estimated effective number of parameters; Δ_IC_ IC_LOO_ difference to best model; SE() standard error. See Table 1 for predictor variable descriptions. The cross validation procedure yields an estimate of the information criterion, and the associated uncertainty. Model comparison is therefore based not just on Δ_IC_ values (and arbitrary difference thresholds), as is routinely done with AIC differences in maximum likelihood frameworks, but by taking the standard error for the estimated Δ_IC_ into account.

| Model | IC<sub>LOO</sub> | SE(IC<sub>LOO</sub>) | $\Delta_{IC}$ (SE) | p<sub>LOO</sub> | SE(p<sub>LOO</sub>) |
| --- | --- | --- | --- | --- | --- |
| TEMP + GOSNER + log(DENS) +<br>WLEV + log(PLANK) + log(DIST) | 704 | 65 |  | 117.4 | 15.4 |
| TEMP + GOSNER + log(DENS) +<br>WLEV + log(PLANK) | 712 | 67 | 8 (18) | 113.2 | 14.7 |
| TEMP + GOSNER + log(DENS) +<br>WLEV + log(DIST) | 989 | 116 | 286 (85) | 115.5 | 22.1 |
| TEMP + GOSNER + log(DENS) +<br>WLEV | 1003 | 112 | 300 (84) | 116.8 | 22.5 |

